# Phylogenetic patterns over sixty-five years of vegetation change across a montane elevation gradient

**DOI:** 10.1101/2025.11.07.687265

**Authors:** Leah N. Veldhuisen, Stephanie D. Zorio, Charles F. Williams, Katrina M. Dlugosch

## Abstract

Phylogenetic diversity is an important axis of biodiversity associated with ecosystem stability and productivity. However, climate change will threaten phylogenetic diversity when extinctions are phylogenetically clustered. Despite the importance of phylogenetic diversity for current and future ecosystems, and the potential unique insights it offers into communities, few studies have quantified its long-term changes. Here, we leverage a dataset spanning approximately 65 years and 1500 m of elevation in the Colorado Rocky Mountains to test for changes in phylogenetic diversity in angiosperm communities. We included four community types (sagebrush, spruce-fir, upland-herb, alpine), which vary in the direction and magnitude of changes in species richness over this period. We tested whether individual species’ responses to climate change could be predicted by phylogeny, including changes to abundance, constancy (% of sites occupied), and elevational range. We did not find phylogenetic signal in abundance change in any community, but we did find phylogenetic signal in both constancy shifts in alpine and elevational shifts in sagebrush communities. We then tested for changes in phylogenetic diversity across time for each community type. We found that phylogenetic diversity changed in the same direction as species richness in the sagebrush community, where both declined, and in the upland-herb, where both increased, with important roles for both species gains and losses in community phylogenetic composition. The alpine community did not change in phylogenetic diversity, although richness declined, while the spruce-fir community increased in phylogenetic diversity but did not change in richness, emphasizing the potential for changes in different aspects of diversity to be decoupled. Our work highlights that the impacts of climate change vary across communities, and that phylogeny is shaping changes in certain systems. Given the importance of phylogenetic diversity to ecosystem function, long term studies are essential for understanding how climate change impacts phylogenetic diversity in distinct community types with unique responses to climate change.

## Introduction

Phylogenetic diversity is a key axis of biodiversity that, in plants, has been associated with ecosystem stability and productivity (Cadotte, Cardinale, and Oakley 2008; Cadotte, Dinnage, and Tilman 2012; Cadotte 2013; Davies et al. 2016; Maherali and Klironomos 2007). Importantly, phylogenetic diversity can capture variation among species that is missed when only studying one or a few traits, which may more accurately reflect the niche space occupied by species and communities (Huang et al. 2020; E-Vojtkó et al. 2023; Cadotte et al. 2009). As a result, phylogenetic diversity cannot necessarily be assumed to correlate with other diversity metrics, such as species richness or functional diversity (Gerhold et al. 2015; E-Vojtkó et al. 2023; Hähn et al. 2024; Veldhuisen et al. 2025a). Research also suggests that climate change threatens phylogenetic diversity as a result of phylogenetically clustered extinction in plant communities (Li, Miller, and Harrison 2019; Thébault, Huber, and Loreau 2007; González-Orozco et al. 2016; Eiserhardt et al. 2015; Willis et al. 2008; Padullés Cubino et al. 2024; López-Rubio et al. 2022). Despite the importance of phylogenetic diversity for ecosystems, and the potential insights it offers into communities, few studies have quantified its long-term changes or the phylogenetic patterns of individual species’ changes (Willis et al. 2008; Li, Miller, and Harrison 2019; Padullés Cubino et al. 2024; Widmer et al. 2025), as appropriate high quality long term datasets are rare (Kapfer et al. 2016). Here, we take advantage of a dataset spanning 65 years and a 1500 m elevational gradient (Zorio, Williams, and Aho 2016; Langenheim 1962) to test for changes in phylogenetic diversity over time and space in Rocky Mountain angiosperm communities.

The Rocky Mountains of North America are experiencing significant impacts of climate change, including increased temperatures, reduced summer rain and winter snow, and earlier snowmelt (Miller-Rushing and Inouye 2009; Mote et al. 2005; Bolinger et al. 2023). Plants in the Colorado region of the Rocky Mountains are particularly well studied, with changes reported in flowering phenology (CaraDonna, Iler, and Inouye 2014; Inouye 2008; Lambert, Miller-Rushing, and Inouye 2010), plant-pollinator interactions (CaraDonna et al. 2017; Bain et al. 2022; Iler et al. 2019; Arrowsmith et al. 2023; Gallagher and Campbell 2017), and plant functional traits (Powers, Briggs, and Campbell 2025; Powers et al. 2022; Stark et al. 2017). Many studies have also documented changes in traditional alpha and beta diversity metrics and in species’ abundance (Zorio, Williams, and Aho 2016; CaraDonna, Iler, and Inouye 2014; Inouye 2008; Rudgers et al. 2014; Langenheim 1962; Arrowsmith et al. 2023; Harte and Shaw 1995; Panetta, Stanton, and Harte 2018). In particular, CaraDonna et al. (2014) found significant changes in flowering phenology over 40 years, and CaraDonna & Inouye (2015) found that these same species exhibited phylogenetic signal in their first and peak flowering dates. These phylogenetic signatures in individual species’ responses to climate change suggest that the phylogenetic composition of communities as a whole may also be changing in this system.

A set of earlier studies in the Colorado Rocky Mountains offers an exceptional opportunity to directly observe how entire communities have changed over more than half a century in this region. In her foundational work in Colorado’s East River Valley, Jean Langenheim quantified plant species cover for four different community types (sagebrush, spruce-fir, upland-herb and alpine) across elevations from 2600 m to 4100 m from 1948 to 1952 (Langenheim 1962; Langenheim 1953). Zorio and colleagues then resampled these same sites in 2012-2014, approximately 65 years after Langenheim’s surveys (Zorio, Williams, and Aho 2016; S. Zorio 2015). Zorio et al. (2016) found distinct changes in species richness, alpha and beta diversity, and species’ median elevations and abundances from circa 1950 to 2014. When pooled across all elevations, species richness increased significantly, although results varied by community type (Zorio, Williams, and Aho 2016). In the upland-herb communities, richness and Shannon diversity increased significantly over this 65-year period, while in alpine communities, both metrics decreased significantly. There were no significant changes in the spruce-fir community, while only richness declined significantly in the sagebrush community (Zorio, Williams, and Aho 2016). Zorio et al. (2016) also found that sites within a single community type became more heterogeneous during this period in all but the alpine zone, and that differences in species composition among community types became less distinct. Community changes were largely driven by declines in the most abundant species in all four community types, as well as changes in growth forms and the extent of bare ground (Zorio et al. 2016). These results reveal that community composition is indeed changing in the region, but that responses differ by community type. Additionally, Zorio et al. (2016) did not investigate phylogenetic composition of these communities.

Here, we use data from Langenheim (1953, 1963) and Zorio et al. (2016) to investigate how phylogenetic diversity has changed in these Rocky Mountain plant communities from circa 1950 to 2014 (Zorio, Veldhuisen, and Williams 2025). We first test for phylogenetic clustering (signal) in individual species’ changes over this period, because correlated responses in closely related species should disproportionately affect the phylogenetic composition of their communities. In this dataset, species responses are available as changes in abundance (% cover), changes in constancy (the % of sites occupied within a community type), and changes in observed mean elevational range across all community types. We test for phylogenetic signal in each of these three metrics. We then test for changes in community-level phylogenetic diversity over time, using multiple metrics of phylogenetic diversity, and ask whether responses differ significantly by community type, as communities have experienced declines (sagebrush, alpine), increases (upland-herb), or no change (spruce-fir) in species richness during this period (Zorio, Williams, and Aho 2016). We ask whether phylogenetic patterns in individual species’ changes and community diversity differ across multiple community types responding to climate change *in situ* over more than six decades.

## Methods

### Study Area and Data Collection

The plant communities originally surveyed by Langenheim (Langenheim 1962; Langenheim 1953) are distributed across a wide elevational gradient (2600 m to 4100 m) in and around the East River Valley near the Rocky Mountain Biological Laboratory in Gothic, CO, USA (38.8697°N, 106.9878°W; Fig. 1). Average annual temperature increased from 36℃ to 38℃ from 1950 to 2014 (NOAA 2014). Average precipitation was 4% lower in the 2001-2022 timeframe than in the 1950-2000 timeframe (Fig. 2B), with a larger 6% decline specifically in average summer precipitation (Bolinger et al. 2023). The region’s relatively short growing season lasts 3-5 months; snowmelt date is variable, but has been earlier on average with climate change (CaraDonna, Iler, and Inouye 2014; Aldridge et al. 2011). As of the second round of data collection in 2012-2014, the average annual maximum temperature in this region was 10.8°C, and the average annual minimum temperature was –7.8°C (Zorio, Williams, and Aho 2016; NOAA 2014), with an average snowfall of 502.7 cm per year (Zorio, Williams, and Aho 2016; NOAA 2014).

**Figure 1:**
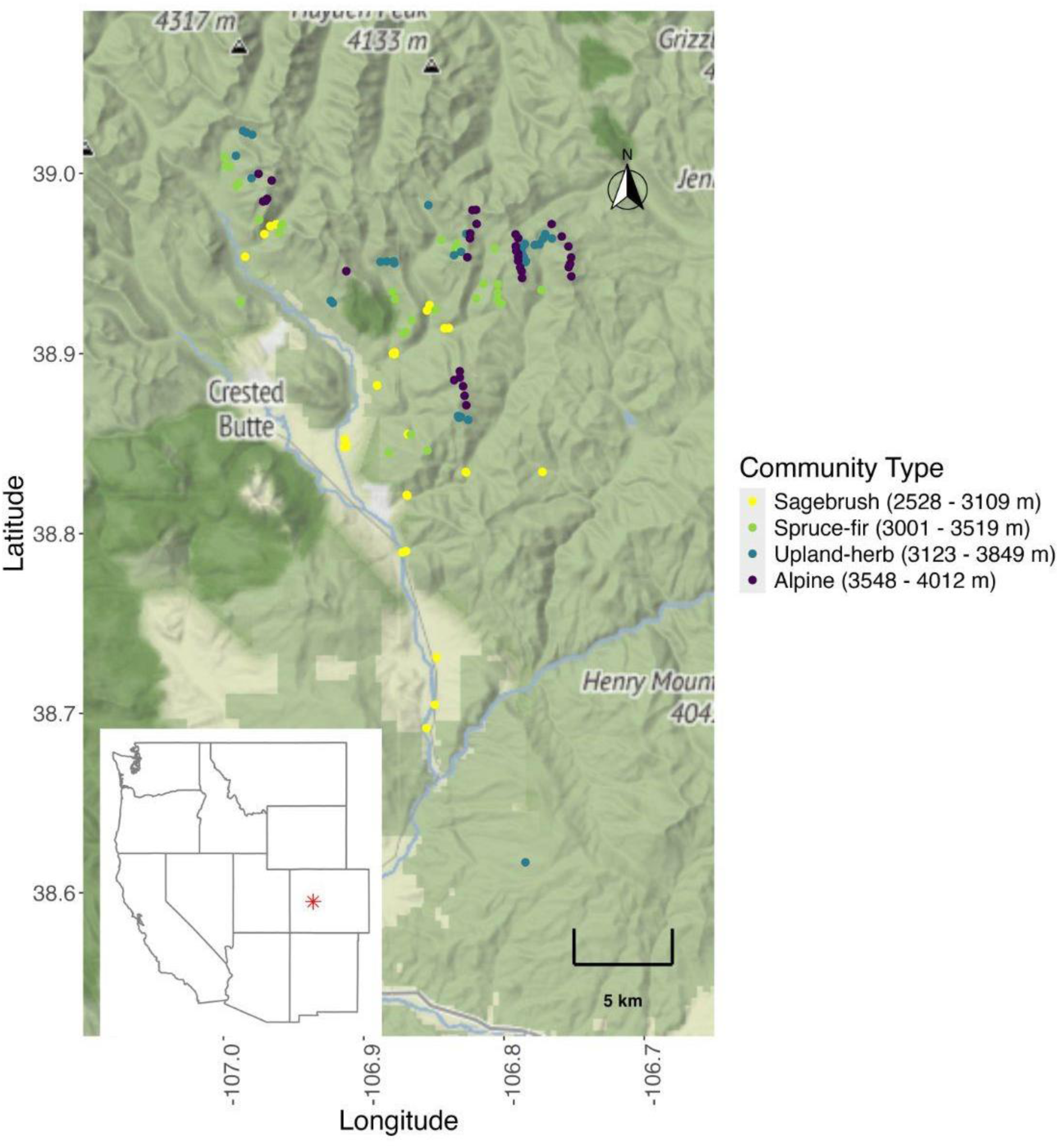
Map of the East River Valley and surrounding area with site locations color-coded by community type (yellow for sagebrush, light green for spruce-fir, blue for upland-herb and purple for alpine). Inset shows the Western United States with the general study location marked with a red star.

**Figure 2:**
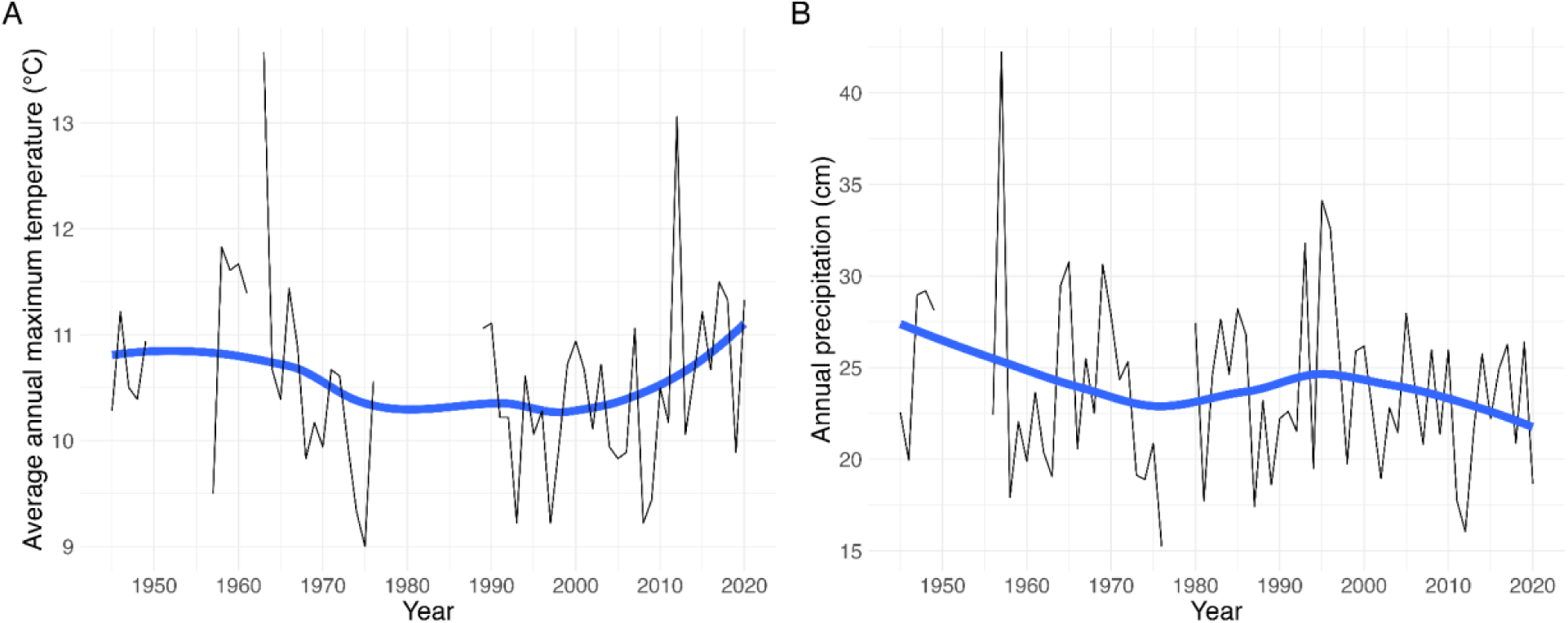
Graphs showing changes in A) average annual maximum temperature (°C) and B) annual precipitation (cm) at the Crested Butte, CO weather station from 1945 to 2020. Blue lines are loess smoothing lines. Data were downloaded from the Colorado Climate Center.

From 1948 to 1952, Jean Langenheim collected quantitative abundance data from 125 sites across four community types for her Ph.D. research (Langenheim 1962; Langenheim 1953). She defined four community types by their dominant species: from lowest to highest elevation, sagebrush (2528 m to 3109 m), spruce-fir (3001 m to 3519 m), upland-herb (3123 m to 3849 m), and alpine (3548 m to 4012 m). Upland-herb communities are now commonly referred to as subalpine meadows; we use ‘upland-herb’ here to maintain consistency with Langenheim’s phrasing (Langenheim 1962; Zorio, Williams, and Aho 2016). Langenheim conducted her surveys with the paced transect method (Levy & Madden, 1933): she recorded each plant that touched her boot at approximately 1-m intervals (Langenheim, 1962). Zorio et al. (2016) used an equivalent method, using a 75 m transect tape and recording intersecting species every meter.

Through discussion with Drs. Jean and Ralph Langenheim, original site descriptions, photos, and herbarium specimens, Zorio and colleagues (2016) relocated and resampled 121 of Langenheim’s 125 sites (27 in sagebrush, 31 in spruce-fir, 30 in upland-herb and 33 in alpine; Fig. 1). A fifth community type, aspen forest, was included in Langenheim (1953), but there were insufficient geographic data available for Zorio et al. (2015, 2016) to relocate those sites. Following Zorio et al. (2016), we do not consider aspen forests here. GPS coordinates, elevation, slope, aspect and substrate for each site are shown in the sheet titled ‘Lang Zorio Env Site Data’ in the data repository (Zorio, Veldhuisen, and Williams 2025).

Notably, Langenheim did not report abundance data for species with less than 14% constancy (i.e., those occurring in fewer than 14% of total sites) in her original dataset, and she grouped some taxa that were difficult to identify to species by genus. Langenheim (1962) stated that grouped taxa mostly included grasses, sedges and *Vaccinium,* although it is likely that some Asteraceae species were also unintentionally grouped (Zorio, Williams, and Aho 2016). To ensure that the 2014 dataset was comparable with the 1950 dataset, Zorio et al. (2016) reported all species but used a reduced and subsampled dataset with only species exhibiting >14% constancy and lumped species as in Langenheim to calculate changes in species richness and Shannon diversity (see Zorio 2015 and Zorio et al. 2016 for more detail). We also use the 14% constancy cutoff for our analyses for consistency and to avoid artificially inflating richness and diversity values in 2014.

### Datasets

We used two datasets for our analyses, both available in Zorio, Veldhuisen, and Williams (2025). The first, which we will refer to as the short dataset, was generated by Zorio et al. (2016) and includes changes in mean elevation, constancy, and abundance for species that were present and identified in both the 1950 and 2014 surveys. It includes elevation shifts for 83 species and abundance and constancy data for 79 species. We used this short dataset to calculate phylogenetic signal for changes in elevation, constancy and abundance (see *Phylogenetic signal* section below). This dataset available in our data repository (Zorio, Veldhuisen, and Williams 2025), under the files labelled “Zorio_abundance_and_constancy_change” and “Zorio_elevation_change.”

The second dataset, which we will refer to as the full site-level dataset, contains site-level presence/absence data for all species at all sites in 1950 and 2014. Langenheim’s entire thesis (Langenheim 1953) with these data is also archived at RMBL. In the 1950 full site-level data with the 14% constancy cutoff, both the sagebrush and spruce-fir communities contained 43 species, the upland-herb community contained 59 and alpine contained 50. For the 2014 full site-level data, we have richness values with and without the 14% constancy cutoff. We report both, with the total number listed first and the 14% cutoff richness in parenthesis. The sagebrush community contained 143 species (86), the spruce-fir community contained 128 species (49), the upland-herb contained 168 species (49), and alpine contained 107 species (47). We use the full site-level dataset with the 14% constancy cutoff for the phylogenetic diversity analyses described in the *Phylogenetic diversity metrics* section below.

### Phylogenetic signal

To test for phylogenetic signal in individual species changes over time, we used the (Smith and Brown 2018) “ALLMB.tre” angiosperm phylogeny and Zorio and colleagues’ (2016) short dataset with elevation, abundance and constancy changes for each species. Smith & Brown’s (2018) phylogeny includes all genera in our dataset. Of the 83 species for which we had abundance and constancy change data, only three were not in the Smith & Brown (2018) phylogeny (not including lumped taxa and updated names). Of 79 species with elevation shift data, only seven were not in the phylogeny (also not including lumped taxa and updated names). We used Zorio et al.’s (2016) Table 2 footnotes to inform our phylogenetic assignments for Langenheim’s lumped taxa. For each genus containing multiple species at a given elevation, we used the first species listed in the footnotes to represent the lumped taxa. If the first listed species was not in Smith & Brown, we included the second listed species. All species listed first or second by Zorio et al. were included in Smith & Brown’s phylogeny, except for *Erigeron*, where it was necessary to use the fourth listed species. Four genera (*Poa, Lupinus, Astragalus, Salix*) in the 1950 data did not have individual species noted in Zorio et al. (2016). For these genera, we checked for herbarium specimens deposited by Jean Langenheim at the relevant elevation from 1948-1952 in regional herbaria (RMBL, COLO, CS). If we could not find an herbarium specimen, we chose a representative species for the community type based on *Flora of Colorado* (Ackerfield 2015). For taxa identified to species level that were not in the Smith & Brown (2018) phylogeny, we chose a closely related congener also known to live in the area. We used updated species names as necessary using the Taxonomic Name Resolution Service (Boyle et al. 2013). For all substitutions and name updates for the short dataset, see our Zenodo repository Table S1 (Veldhuisen 2025).

We calculated phylogenetic signal for changes in local abundance, constancy and median elevation using Blomberg’s K (Blomberg et al., 2003) and Pagel’s λ (Pagel, 1999). Zorio and colleagues’ (2016) elevation shift data show the difference in each species’ average elevation from 1950 to 2014. As such, we calculated phylogenetic signal in elevation shift for all communities combined. We also split species into community type and calculated signal in elevation shift by community, though species did not have unique elevation shift values for different communities. For abundance and constancy changes, we calculated phylogenetic signal for each community type. We calculated Blomberg’s K with the “phylosignal()” function in the ‘picante’ R package (Kembel et al., 2010) with 5000 randomizations, and Pagel’s λ with the function “phylosig()” in the ‘phytools’ R package (Revell, 2011) with 5000 iterations and model set to “lambda.” We calculated phylogenetic signal two different ways: once with the described species substitutions, and once removing all genera with lumped species. Blomberg’s K or Pagel’s λ values near 1 indicate trait evolution consistent with Brownian motion. In contrast, K less than 1 or Pagel’s λ near zero suggest less phylogenetic signal than expected by Brownian motion.

### Phylogenetic diversity metrics

To test for changes in community-level phylogenetic diversity across time, we again used the Smith and Brown (2018) “ALLMB.tre” phylogeny and the full dataset with site-level community composition. We used the same methods as described above to determine representative species and to replace species missing from Smith & Brown (2018). Specifically, for the 2014 full site-level data, if the genus in question did not have a footnote in Zorio et al.’s (2016) Table 2, we checked for herbarium specimens by the original authors (Stephanie Zorio, Charles F. Williams, Jean Langenheim) from the year of data collection. Some 2014 specimens had notes stating that they were collected during resampling Langenheim sites, and we used these to inform substitutions in the 2014 full site-level dataset if available. For some genera in Asteraceae with many congeners, we found no phylogenies to inform replacements and many of the species are missing from the Smith and Brown phylogeny. Consequently, we could not include 12 Asteraceae species. For all substitutions and name updates for the full site-level datasets, see our Zenodo repository Table S2 (Veldhuisen 2025). As for phylogenetic signal, we also calculated phylogenetic diversity two different ways: once with the described species substitutions and once simply removing all genera with lumped species.

For metrics of community phylogenetic diversity, we calculated Faith’s phylogenetic diversity (PD; (Faith, 1992)), mean phylogenetic distance (MPD), and mean nearest taxon distance (MNTD) and their associated standard effect sizes (SES) in ‘picante’ (Kembel et al., 2010). Faith’s PD (Faith, 1992) is the total evolutionary distance (branch length) across a phylogenetic tree describing the relationships among the taxa in a dataset. PD will increase if relationships are more distant across the tree and species are more evolutionarily divergent. MPD is the mean distance between pairs of species. Like PD, MPD also increases as evolutionary relationships are more distant across the tree. However, it will be less sensitive to rare cases of distantly-related species that would otherwise elevate Faith’s PD. MNTD is the mean of the distance to the closest relative of each taxon, and reflects the typical distance to any species’ closest relative in the dataset and increases as closest relatives are more distant. MNTD is more indicative of the tendency of species to have closer relatives than it is of the overall evolutionary diversity across the entire tree (community of species) that is captured by the former metrics (for further discussion, see review by (Tucker et al., 2017)).

To calculate SES, we used each metric’s “sample.pool” null model to randomly draw species from the regional species pool with 5000 iterations. We defined our species pool as all species observed in all community types at both time points. We included species below the 14% constancy cutoff in our regional species pool, since they live in the area and have the chance to occur at higher abundances. Positive SES values indicate that the species group in question is more phylogenetically diverse than would be a random draw of the same number of species from the regional species pool (overdispersion). SES values above 1.96 will have P values above 0.975, indicating the SES’s statistical significance. Negative SES values indicate that the species group in question is less phylogenetically diverse than a random draw of the same number of species from the regional species pool (underdispersion). SES values below -1.96 will generate P values below 0.025, indicating the SES’s statistical significance.

### Analysis of phylogenetic diversity metrics

To test for changes in phylogenetic diversity, we used linear models. First, we subtracted the 2014 SES values for each metric and site from the 1950 SES for the same metric and site to get ΔSES values for each site. We included site elevation, community type and slope aspect as explanatory variables. We used ΔSES as our response variable in the model, and used a sum to zero contrast for community type and slope aspect and scaled elevation. We fit separate models for each phylogenetic diversity metric (PD, MPD, MNTD). We also fit separate models for data including lumped taxa with the corresponding substitutions and with the datasets where we removed all lumped taxa.

## Results

### Community phylogenies

The full phylogenetic tree, including all replacements and representatives for lumped taxa, included 318 species across all community types and both 1950 and 2014 data collection periods (Appendix S1: Figure S1). We retained species below the 14% constancy cutoff in our regional pool to calculate standard effect sizes for phylogenetic diversity metrics, therefore they are counted in the total 318 species. The short dataset that we used to calculate phylogenetic signal in elevation shift contained all 83 species in its phylogenetic tree; similarly, the short dataset with abundance and constancy data contained all 79 species in the dataset in its tree. Once we removed species without phylogenetic data, the 1950 sagebrush community phylogenetic tree contained 43 species, the spruce-fir community contained 41, the upland-herb community contained 54, and the alpine community contained 49 species. For the 2014 full site-level data, the sagebrush community contained 86 species, the spruce-fir community contained 49 species, the upland-herb contained 49 species, and alpine contained 47 species in their phylogenetic trees.

### Phylogenetic signal

Removing lumped taxa made no difference to the statistical significance of any of our tests for phylogenetic signal in changes over time, so here we report only tests that included lumped taxa using their representative species (Veldhuisen 2025: Table S2). Results for all calculations with and without lumped taxa are in our Zenodo repository Table S3 (Veldhuisen 2025). For elevation shift, we found significant phylogenetic signal for the sagebrush community with Blomberg’s K (Blomberg, Garland, and Ives 2003) (Fig. 3A; K = 0.29, *p* = 0.04). This effect was largely driven by decreasing average elevation in Asteraceae species (Fig. 3A). No other community types, nor all species combined, showed significant phylogenetic signal in elevation shifts (all *p* > 0.05). For constancy shifts, we found significant phylogenetic signal for the alpine community by both metrics (Fig. 3B; K = 0.53, *p* = 0.01, λ = 0.68, *p* = 0.001), but no significant phylogenetic signal in constancy changes for any other community type (all *p* > 0.05). The species that drove the signal in alpine constancy change were members of the Fabaceae family, which increased in constancy from 1950 to 2014 (Fig. 3B). For abundance shifts, we found no significant phylogenetic signal for any community type for either Blomberg’s K (Blomberg, Garland, and Ives 2003) or Pagel’s λ (Pagel 1999) (Fig. 4; all *p* > 0.05).

**Figure 3:**
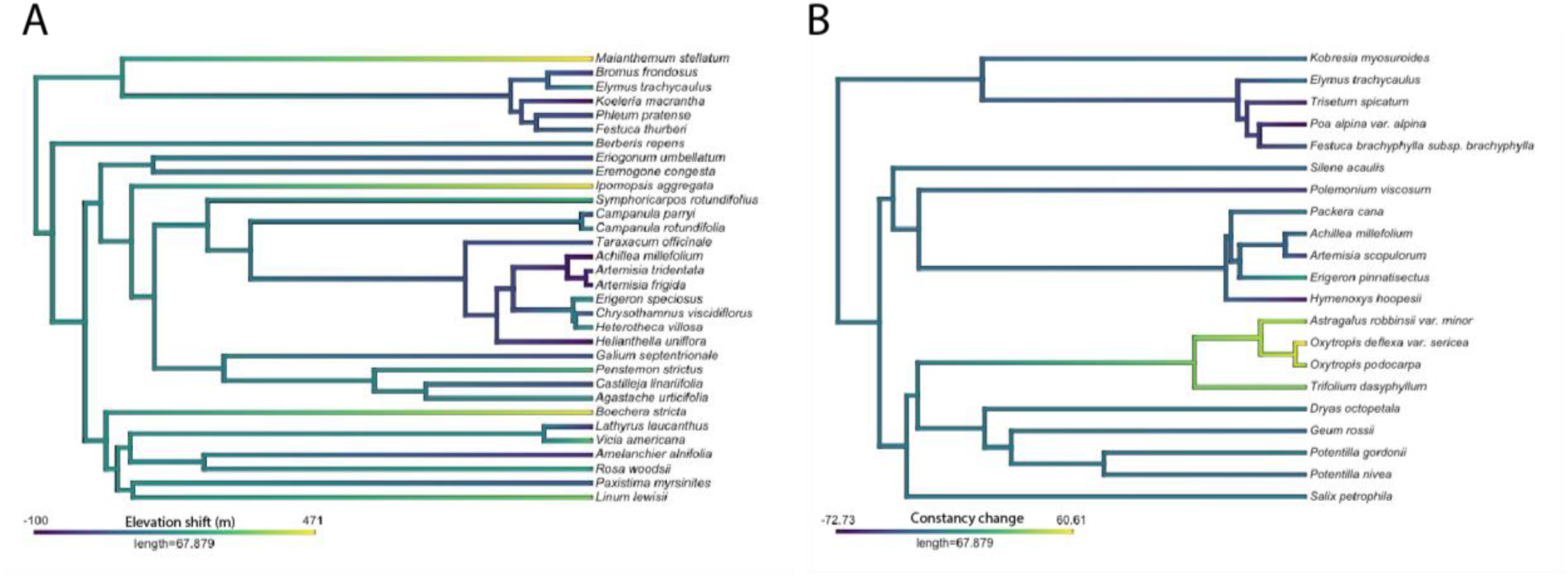
Phylogenies showing significant phylogenetic signal in A) elevation shift in the sagebrush community (K = 0.29, *p* = 0.04,) and B) constancy change in the alpine community (K = 0.53, *p* = 0.01; λ = 0.68, *p* = 0.001). Darker purple shows decreasing average elevation or constancy, while warmer green and yellow shows increasing elevation or constancy.

**Figure 4:**
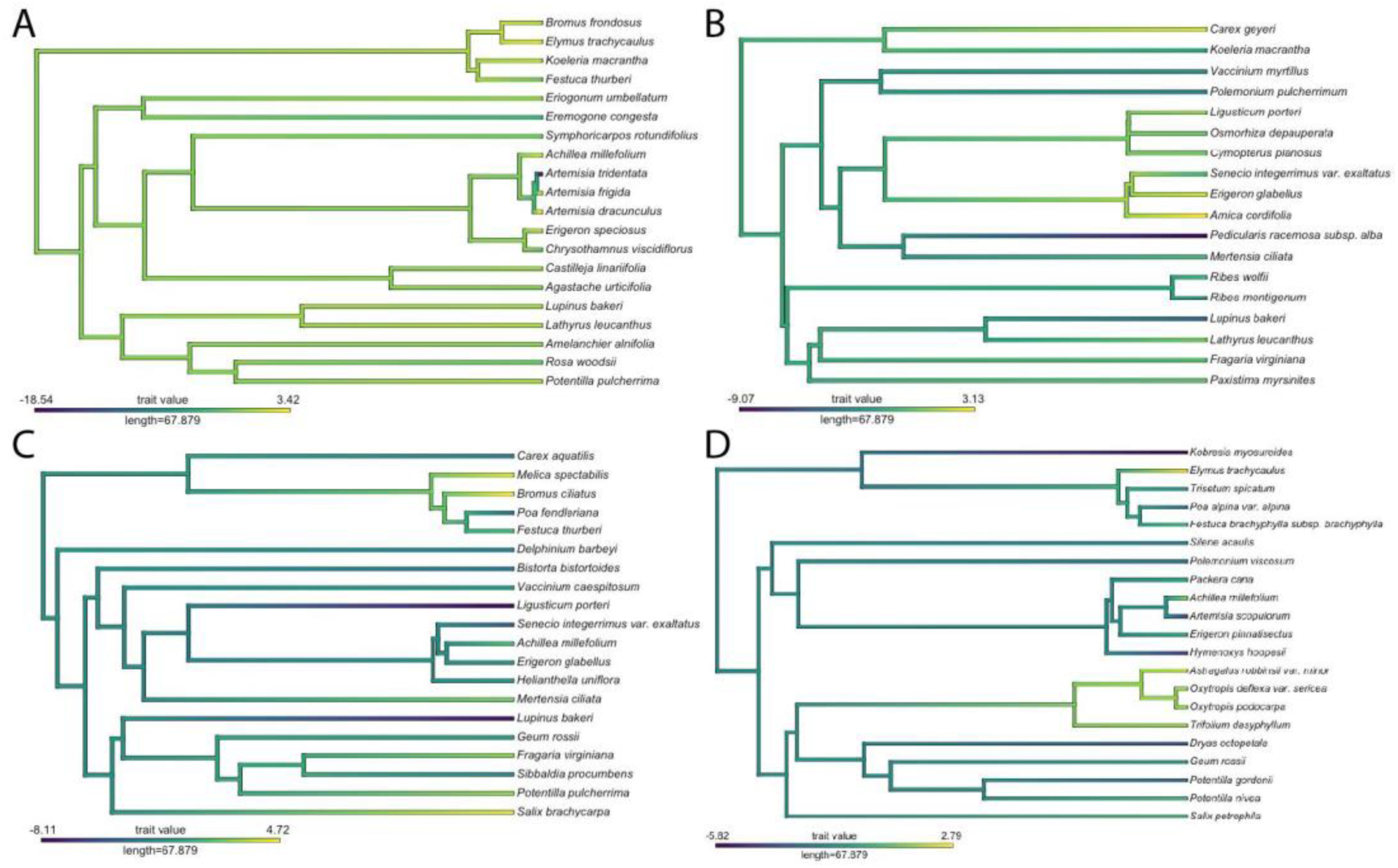
Phylogenies showing no phylogenetic signal for abundance change any community (A: sagebrush, B: spruce-fir, C: upland-herb, D: alpine). Darker purple shows decreasing abundance, while warmer green and yellow shows increasing abundance.

### Change in phylogenetic diversity

Community type was a significant predictor for change in all three diversity metrics (all *p* < 0.01). An effect of site elevation was not significant in any case (all *p* > 0.05), and neither was slope aspect (all *p* > 0.05). With the inclusion of lumped taxa, the sagebrush community declined in phylogenetic diversity, spruce-fir and upland-herb increased, and alpine did not change, with changes being generally consistent across metrics of diversity (Fig. 5). In the sagebrush community, PD (*p* = 0.007), MPD (*p =* 0.048), and MNTD (*p* = 0.004) all declined. In the spruce-fir community, PD did not change (*p* = 0.128), but both MPD (*p* = 0.004) and MNTD (*p* = 0.012) increased. In the upland-herb community, PD (*p* < 0.01) and MNTD (*p* < 0.01) increased, while MPD also increased but not significantly (*p* = 0.07). In the alpine community, none of the three metrics changed significantly (all *p* > 0.05).

**Figure 5:**
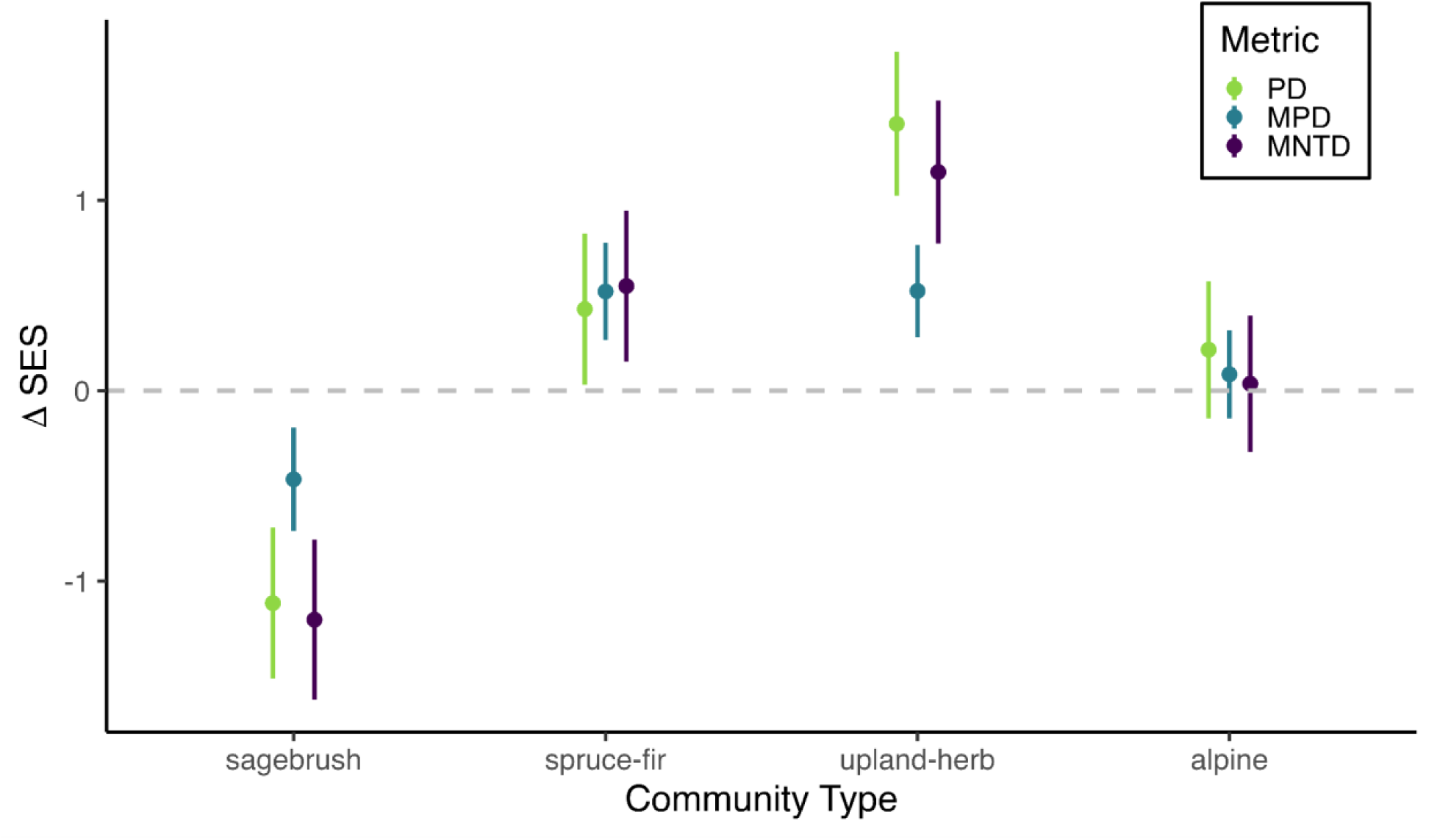
Changes in standard effect size for Faith’s phylogenetic diversity (green), mean phylogenetic distance (blue) and mean nearest taxon distance (purple) for each community type. Points represent means ± SE. Change is statistically significant if the error bars do not cross the grey dashed line, which includes all changes in sagebrush and upland-herb, none in alpine, and MPD and MNTD (but not PD) in spruce-fir.

When we removed lumped taxa, our results changed for the sagebrush and spruce-fir communities, but not for upland-herb or alpine communities (Appendix S1: Figure S2). We removed 21 taxa from the sagebrush community, 12 taxa from spruce-fir, 22 taxa from upland-herb and 12 taxa from alpine. Without lumped taxa, the significant decline in all three phylogenetic diversity metrics in the sagebrush community disappeared (all *p* > 0.05). In the spruce-fir community, the significant increases in MPD and MNTD became non-significant (both *p* > 0.05), and the change in PD stayed non-significant (*p* > 0.05. The significant increase in upland-herb across all three metrics remained the same (PD: *p* < 0.01; MPD: *p* = 0.03; MNTD: *p* < 0.01), while alpine continued to show no significant changes in any metric (all *p* > 0.05).

## Discussion

We used a long-term dataset that spans 65 years and a 1500 m elevation gradient to test whether climate change has affected the phylogenetic diversity of Rocky Mountain plant communities. We examined changes in four community types known to differ in whether their species richness had increased, decreased, or remained unchanged during this time (Zorio, Williams, and Aho 2016). We first tested for phylogenetic signal (clustering) in individual species changes over time within each community, in terms of their abundance within sites, constancy across sites, and elevational range. We found evidence for phylogenetic signal in elevational change and constancy change, but only in one community type for each metric. We then tested for changes in phylogenetic diversity metrics over time in each community. We found that changes in phylogenetic diversity were variable across community types, and that phylogenetic diversity changes were sometimes, but not always, in the same direction as changes in species richness. Further, our results suggest that individual species’ responses are, for the most part, not strongly phylogenetically conserved, highlighting that many species will respond to climate change differently than their closest relatives in the same community.

Overall, we found significant changes in phylogenetic diversity in three of four community types, including both declines and increases. In two community types, the decline or increase in phylogenetic diversity aligned with changes in richness, for the low elevation sagebrush and mid-high elevation upland-herb, respectively. Declines in richness in the alpine were phylogenetically distributed and resulted in no change in phylogenetic diversity, whereas non-significant changes in richness in the spruce-fir community resulted in increased phylogenetic distance among community members. These results align with previous work indicating that impacts of climate change on plant communities are not homogenous across elevations or community types (Crimmins, Crimmins, and David Bertelsen 2009; Harrison, Damschen, and Grace 2010; Palmquist et al. 2021; García Criado et al. 2025; Aldridge et al. 2011; Widmer et al. 2025). We are among the first studies to address changes in phylogenetic diversity over decades (Willis et al. 2008; Li, Miller, and Harrison 2019; Padullés Cubino et al. 2024; Widmer et al. 2025), and the first to do so in the Rocky Mountains. Broad patterns remain under investigation, but we identify some emerging patterns. First, we find evidence that the community’s position within elevation gradient matters, as warmer and drier versus colder and wetter community types experienced declines versus increases in phylogenetic diversity, respectively. Second, certain communities experienced more colonization over the 65 year timespan than others (Zorio, Williams, and Aho 2016), and the phylogenetic distribution of colonizing species significantly impacts phylogenetic diversity. Whether newcomers are invaders or natives shifting ranges, their phylogenetic relationship to the rest of the community is key to existing phylogenetic diversity. Continued work to elucidate these two patterns is essential for pinpointing where climate change poses the biggest threat to native plant diversity and ecosystem functioning.

Our findings are similar to those from a six-year experimental warming study in the same region: both studies suggest that phylogenetic diversity can be coupled or decoupled from traditional metrics of community diversity, indicating that it provides unique information about a community’s response to change (Veldhuisen, et al. 2025a). Veldhuisen et al. (2025a) found that six years after transplanting entire plant communities up and down an elevation gradient to simulate climate change, species richness and Shannon diversity generally increased with experimental warming. This finding is consistent with Zorio et al.’s (2016) observation of natural changes over time. Veldhuisen et al. (2025a) also found that changes in phylogenetic diversity did not follow patterns in richness and Shannon diversity, which contrasts with our findings in the sagebrush and upland-herb communities, but aligns with our findings in the spruce-fir and alpine communities. The authors also tested for phylogenetic signal in abundance changes of individual species in response to these treatments (i.e., whether more closely related species responded similarly, which would be expected if there is phylogenetically clustered extinction risk; (Eiserhardt et al. 2015; Li, Miller, and Harrison 2019; Willis et al. 2008). They did not identify significant phylogenetic signal in abundance response to any treatment (Veldhuisen et al. 2025a), again aligning with our findings of no phylogenetic signal in abundance change over time.

The communities dominated by sagebrush (*Artemisia tridentata* Nutt.) occur at the lowest elevations in our study, in areas that are the warmest and driest among our four community types (Ackerfield 2015). Sagebrush communities experience less snow during the winter and are covered by snow for shorter periods than higher community types (Sloat et al. 2015). We found evidence of declines across all phylogenetic diversity metrics, which parallels Zorio et al.’s (2016) observed declines in richness and Shannon diversity in these sites. These results are in line with previous work suggesting that more water-limited sites may experience bigger declines in diversity with warming (Kelly and Goulden 2008; Harrison, Damschen, and Grace 2010). Similarly, a recent modeling study across the western US predicted that climate change would drive significant changes in *A. tridentata* populations and associated communities (Palmquist et al. 2021). Another recent study in central Europe found that both functional and phylogenetic diversity declined most at the lowest elevation sites from the late 1800s to current day (Widmer et al. 2025). Widmer et al. (2025) posit that their low elevation diversity declines were driven by land use change and agriculture during the Third Agricultural Revolution; land use change would also be an important future avenue in our study sites, as it is likely also more intense at our lower elevation sites. Zorio et al. (2016) noted that the sagebrush community also experienced increased invasion, particularly by the grass *Bromus inermis* Leyss. While we know that *B. inermis* is closely related to other native grasses in the community, we cannot disentangle if invasion decreased phylogenetic diversity (Daru et al. 2021; Selvi, Carrari, and Coppi 2016) or if lowered diversity facilitated invasion (Galland et al. 2019; Ketola, Saarinen, and Lindström 2017). This would again be an important avenue for future work. Zorio et al. (2016) also reported that species in sagebrush had the greatest upwards elevation shifts over time, with many species shifting into new community types. Notably, we found significant phylogenetic signal in elevation shift for the sagebrush community, but this phylogenetic pattern was largely driven by species in the Asteraceae contracting to lower elevations while most other taxa shifted higher. Combined, these findings suggest that phylogenetically clustered elevation shifts (resulting in richness declines), coupled with invasion by closely related species, are driving declines in overall phylogenetic diversity in this low elevation community type.

At the other extreme are the alpine communities, the highest elevation community type, where Zorio et al. (2016) also found significant declines in both richness and Shannon diversity. Here we found no significant changes in any phylogenetic metric. Of note, the alpine experienced the most significant drop in richness and had the lowest richness in the 2014 data (Zorio et al. 2016), which may have limited our statistical power more than in other community types. The lack of change in phylogenetic diversity indicates that the losses indicated by Zorio el al.’s (2016) declining richness are distributed across the phylogeny and not phylogenetically clustered. A similar pattern of phylogenetically distributed losses has been seen in European forest understory plant communities (Padullés Cubino et al. 2024). The phylogenetic dispersion of species losses also aligns with our finding that neither abundance change nor elevation changes showed phylogenetic signal in alpine communities. Interestingly, the alpine was the only community type whose composition became more heterogeneous between sites (Zorio, Williams, and Aho 2016). In a recent study in the Arctic, researchers found that colder sites had more gains and losses over time than warmer sites (García Criado et al. 2025). Our findings are contrary to expectations, and suggest that this heterogeneity is not phylogenetically structured and does not impact phylogenetic diversity, unlike the changes occurring at the warmer sagebrush sites. Interestingly, a recent study of phylogenetic patterns across the flowering season in these communities found that communities flowering in the warmest (late) season were also the most phylogenetically structured (Veldhuisen, Enquist, and Dlugosch 2023), suggesting that responses to warm extremes might be more phylogenetically conserved than responses to cold.

The next highest elevation sites are those in the upland-herb communities. Here, Zorio et al. (2016) found significant increases in both richness and Shannon diversity, as did Veldhuisen et al. (2025a) in the warmed treatments in the six year transplant study within this community type. We found that phylogenetic diversity also increased in these sites, indicating that the species joining the community over time were more phylogenetically distant and diverse than species leaving the community. In contrast, the transplant study did not show a significant change in phylogenetic diversity over its shorter duration (Veldhuisen et al. 2025a). Other studies have found that both colonizations and extinctions are important contributors to phylogenetic diversity changes (Bennett, Stotz, and Cahill 2014; Winter et al. 2009; Daru et al. 2021). Importantly, Zorio et al. (2016) noted that the upland-herb community had the highest number of species entering and exiting of all community types, likely because it contains more diverse microhabitats across a wider elevational range. It may also be that, similar to the recent study of arctic plants (García Criado et al. 2025) and the results from the alpine communities in this study, colder sites tend to have more gains and losses over time than warmer sites. We also did not find phylogenetic signal for shifts in abundance, constancy, or elevation of existing species in this community type, again suggesting that newcomers were responsible for increasing phylogenetic diversity.

Finally, in the spruce-fir community, we found increasing phylogenetic diversity with two of three metrics (MPD and MNTD), whereas Zorio et al. (2016) reported declines in richness and Shannon diversity that were not significant. Our results suggest that these slight changes in composition must have increased the phylogenetic distances between species, such as through the loss of close relatives. The lack of statistically significant changes in richness, Shannon diversity and PD suggest that overall community composition changed very little, though it was sufficient to alter aspects of the distribution of phylogenetic diversity in the community represented by MPD and MNTD.

Our work adds to the small but growing body of work addressing the critical question of how climate change impacts phylogenetic diversity patterns (Padullés Cubino et al. 2024; Daru et al. 2021; Winter et al. 2009; Veldhuisen et al. 2025a). Other studies to date have found evidence of phylogenetic homogenization (Padullés Cubino et al. 2024; Daru et al. 2021; Winter et al. 2009; Widmer et al. 2025), and that changes cannot be assumed to correlate with other diversity metrics, such as species richness or functional diversity (Gerhold et al. 2015; E-Vojtkó et al. 2023; Hähn et al. 2024; Veldhuisen et al. 2025a). Our results align with this previous work showing that phylogenetic diversity often responds differently than other diversity metrics. A recent study of European forest understory plants found that phylogenetic diversity did not change over a 40 year period, and that lost and gained species were randomly dispersed across the phylogeny (Padullés Cubino et al. 2024). In contrast, Li and colleagues (2019) found that grassland communities lost phylogenetic diversity over a 19-year period due to lower winter precipitation, but only at the local scale. (De Pauw et al. 2021) found that functional and phylogenetic diversity can respond in opposite directions to the same environmental variable, and several other studies have noted the decoupling of these aspects of diversity (García Criado et al. 2025; De Pauw et al. 2021; Winter et al. 2009; López-Rubio et al. 2022; Veldhuisen et al. 2025a). In contrast, Li, Miller, and Harrison (2019) and Miller et al. (2019) found that phylogenetic and functional diversity both declined in their common study system over the same time frame. Given the ecosystem benefits of phylogenetic diversity and its independence of other diversity metrics, long term studies like ours are critical to better understand how it responds to climate change over time under different community contexts (White et al. 2023).

We provide new evidence of climate change’s impacts on phylogenetic diversity over a relatively long time period, although our study also reveals some limitations of our dataset that underscore the challenges of analyzing long term surveys (Kapfer et al. 2016). The lumping of some taxa in Langenheim’s 1950 data impacted our results. For some genera, we were able to infer how many species were lumped together using Zorio et al.’s (2016) Table 2 footnotes and herbarium specimens. For genera not included in Zorio et al.’s (2016) Table 2, we were unable to determine how many species we may have missed in the 1950 data. When we removed lumped taxa from our analyses, the phylogenetic diversity declines in the sagebrush community and increases in the spruce-fir community disappeared, while the increases in upland-herb and no change in alpine were consistent. The sagebrush community had the most lumped taxa to be removed, and their effects on results in this community (and in the spruce-fir) indicate that these taxa were likely significant drivers of change.

The second limitation of this dataset is the exclusion of rare species occurring below 14% constancy in Langenheim’s 1950 data. Although rare species play important roles in ecosystems (Bracken and Low 2012; Mouillot et al. 2013; Leitão et al. 2016), previous work in this system suggests that rare species’ contributions to phylogenetic diversity are similar to those of more common species (Veldhuisen et al. 2025b). This previous work indicates that the exclusion of the rarest species should not have strongly influenced our results, but it is important to note. Additionally, we had to remove some species within Asteraceae, particularly in the *Erigeron* and *Senecio* genera, that did not occur in the Smith & Brown (2018) phylogeny and had no phylogenetic information or herbarium specimens to inform substitutions. *Erigeron* and *Senecio* are common in the region, and removing them could have altered our phylogenetic diversity results. It would be especially useful for future work to investigate the taxonomy of endemic Rocky Mountain Asteraceae genera so that all species can be included in phylogenetic analyses of these generally well-studied communities, particularly as these species are highly relevant to community and ecosystem functioning (Bain et al. 2022; Veldhuisen, Enquist, and Dlugosch 2023).

Lastly, this dataset includes only two timepoints. Over ten years of accelerating climate change has passed since Zorio et al.’s 2014 resurvey, and a third survey of these sites would permit further investigation of changes over time in the region. Additionally, it would be ideal to investigate the impacts of environmental variables beyond temperature and precipitation on the trajectories of change across community types. Zorio et al. (2016) speculated about the impact of historical logging, fire and livestock grazing, but additional historical data would be essential to assess these possibilities.

In conclusion, our study of 65 years of community change in the Rocky Mountains revealed unique trajectories for each community type, highlighting two emergent patterns. First, phylogenetic diversity responses to climate change are nuanced and community-specific, as lower, warmer and drier communities lost more phylogenetic diversity over time than higher, cooler and wetter communities. The stronger impact of climate change in more water-limited times and places is consistent with other work in the region (Palmquist et al. 2021; Veldhuisen, Enquist, and Dlugosch 2023), but deserves significantly more research. Second, we find that the phylogenetic distribution of new species determines phylogenetic diversity, and communities with higher turnover may experience different changes than communities with lower turnover. Changes in phylogenetic diversity were aligned with changes in species richness in two community types, revealing the nature of continued community change over longer timescales. In two other communities, changes in only one or the other of phylogenetic diversity or species richness were significant, emphasizing the potential for changes in different aspects of diversity to be decoupled, and the importance of both species gains and losses to community phylogenetic composition. Given the importance of phylogenetic diversity to ecosystem function (Cadotte, Cardinale, and Oakley 2008; Cadotte, Dinnage, and Tilman 2012; Cadotte 2013; Davies et al. 2016; Owen et al. 2019; Molina-Venegas, Ramos-Gutiérrez, and Moreno-Saiz 2020), more research and more long term studies are essential for understanding specifically where and when the impacts of climate change will pose the biggest dangers to native plant communities.

## Acknowledgements

Most importantly, we thank Jean Langenheim for collecting her original dataset. We thank members of the Dlugosch lab, ECOL596A students, and J. Bronstein for manuscript feedback and discussion, D. Park for discussion regarding phylogenetic analyses, and V. Milici for discussion regarding statistical analyses. We also thank A. Johnson and G. Glynn for their assistance with the resurvey of Langenheim’s sites. This work was funded by Rocky Mountain Biological Laboratory, Botanical Society of America, and University of Arizona Graduate & Professional Student Council awards to L.N.V. and United States National Science Foundation award #1750280 and United States Department of Agriculture National Institute of Food and Agriculture award #2023-67013-40169 to K.M.D.

## Author Contributions

L.N.V. and K.M.D. conceived the study. C.F.W. and S.D.Z collected the 2014 field data, and curated the 1950 Langenheim data. L.N.V. analyzed the data. L.N.V. wrote the original draft with input from all authors. All authors provided feedback. All authors approved the final version of the manuscript.

## References

Ackerfield, Jennifer. 2015. Flora of Colorado. Botanical Research Institute of Texas.

Aldridge, George, David W. Inouye, Jessica R. K. Forrest, William A. Barr, and Abraham J. Miller-Rushing. 2011. “Emergence of a Mid-Season Period of Low Floral Resources in a Montane Meadow Ecosystem Associated with Climate Change.” The Journal of Ecology 99 (4): 905–13. 10.1111/j.1365-2745.2011.01826.x.

Arrowsmith, K. C., Victoria A. Reynolds, Heather M. Briggs, and Berry J. Brosi. 2023. “Community Context Mediates Effects of Pollinator Loss on Seed Production.” Ecosphere 14 (6): e4569. 10.1002/ecs2.4569.

Bain, Justin A., Rachel G. Dickson, Andrea M. Gruver, and Paul J. CaraDonna. 2022. “Removing Flowers of a Generalist Plant Changes Pollinator Visitation, Composition, and Interaction Network Structure.” Ecosphere 13 (7): e4154. 10.1002/ecs2.4154.

Bennett, Jonathan A., Gisela C. Stotz, and James F. Cahill Jr. 2014. “Patterns of Phylogenetic Diversity Are Linked to Invasion Impacts, Not Invasion Resistance, in a Native Grassland.” Journal of Vegetation Science: Official Organ of the International Association for Vegetation Science 25 (6): 1315–26. 10.1111/jvs.12199.

Blomberg, Simon P., Theodore Garland, and Anthony R. Ives. 2003. “Testing for Phylogenetic Signal in Comparative Data: Behavioral Traits Are More Labile.” Evolution 57 (4): 717–45. 10.1111/j.0014-3820.2003.tb00285.x.

Bolinger, R. A., J. J. Lukas, R. S. Schumacher, and P. E. Goble. 2023. “Climate Change in Colorado, 3rd Edition.” Colorado State University.

Boyle, B., N. Hopkins, Z. Lu, J. A. R. Garay, D. Mozzherin, T. Rees, N. Matasci, et al. 2013. “The Taxonomic Name Resolution Service: An Online Tool for Automated Standardization of Plant Names.” BMC Bioinformatics 14 (1): 16. 10.1186/1471-2105-14-16.

Bracken, Matthew E. S., and Natalie H. N. Low. 2012. “Realistic Losses of Rare Species Disproportionately Impact Higher Trophic Levels: Loss of Rare ‘cornerstone’ Species.” Ecology Letters 15 (5): 461–67. 10.1111/j.1461-0248.2012.01758.x.

Cadotte, Marc W. 2013. “Experimental Evidence That Evolutionarily Diverse Assemblages Result in Higher Productivity.” Proceedings of the National Academy of Sciences of the United States of America 110 (22): 8996–9000. 10.1073/pnas.1301685110.

Cadotte, Marc W., Bradley J. Cardinale, and Todd H. Oakley. 2008. “Evolutionary History and the Effect of Biodiversity on Plant Productivity.” Proceedings of the National Academy of Sciences of the United States of America 105 (44): 17012–17. 10.1073/pnas.0805962105.

Cadotte, Marc W., Jeannine Cavender-Bares, David Tilman, and Todd H. Oakley. 2009. “Using Phylogenetic, Functional and Trait Diversity to Understand Patterns of Plant Community Productivity.” PloS One 4 (5): e5695. 10.1371/journal.pone.0005695.

Cadotte, Marc W., Russell Dinnage, and David Tilman. 2012. “Phylogenetic Diversity Promotes Ecosystem Stability.” Ecology 93 (sp8): S223–33. 10.1890/11-0426.1.

CaraDonna, Paul J., Amy M. Iler, and David W. Inouye. 2014. “Shifts in Flowering Phenology Reshape a Subalpine Plant Community.” Proceedings of the National Academy of Sciences of the United States of America 111 (13): 4916–21. 10.1073/pnas.1323073111.

CaraDonna, Paul J., William K. Petry, Ross M. Brennan, James L. Cunningham, Judith L. Bronstein, Nickolas M. Waser, and Nathan J. Sanders. 2017. “Interaction Rewiring and the Rapid Turnover of Plant–pollinator Networks.” Ecology Letters 20 (3): 385–94. 10.1111/ele.12740.

Crimmins, Theresa M., Michael A. Crimmins, and C. David Bertelsen. 2009. “Flowering Range Changes across an Elevation Gradient in Response to Warming Summer Temperatures.” Global Change Biology 15 (5): 1141–52. 10.1111/j.1365-2486.2008.01831.x.

Daru, Barnabas H., T. Jonathan Davies, Charles G. Willis, Emily K. Meineke, Argo Ronk, Martin Zobel, Meelis Pärtel, Alexandre Antonelli, and Charles C. Davis. 2021. “Widespread Homogenization of Plant Communities in the Anthropocene.” Nature Communications 12 (1): 6983. 10.1038/s41467-021-27186-8.

Davies, T. Jonathan, Mark C. Urban, Bronwyn Rayfield, Marc W. Cadotte, and Pedro R. Peres-Neto. 2016. “Deconstructing the Relationships between Phylogenetic Diversity and Ecology: A Case Study on Ecosystem Functioning.” Ecology 97 (9): 2212–22. 10.1002/ecy.1507.

De Pauw, Karen, Camille Meeussen, Sanne Govaert, Pieter Sanczuk, Thomas Vanneste, Markus Bernhardt-Römermann, Kurt Bollmann, et al. 2021. “Taxonomic, Phylogenetic and Functional Diversity of Understorey Plants Respond Differently to Environmental Conditions in European Forest Edges.” The Journal of Ecology 109 (7): 2629–48. 10.1111/1365-2745.13671.

Eiserhardt, Wolf L., Finn Borchsenius, Christoffer M. Plum, Alejandro Ordonez, and Jens-Christian Svenning. 2015. “Climate-Driven Extinctions Shape the Phylogenetic Structure of Temperate Tree Floras.” Ecology Letters 18 (3): 263–72. 10.1111/ele.12409.

E-Vojtkó, Anna, Francesco de Bello, Zdeňka Lososová, and Lars Götzenberger. 2023. “Phylogenetic Diversity Is a Weak Proxy for Functional Diversity but They Are Complementary in Explaining Community Assembly Patterns in Temperate Vegetation.” The Journal of Ecology 11 (10): 2218–30. 10.1111/1365-2745.14171.

Gallagher, M. Kate, and Diane R. Campbell. 2017. “Shifts in Water Availability Mediate Plant-Pollinator Interactions.” The New Phytologist 215 (2): 792–802. 10.1111/nph.14602.

Galland, Thomas, Guillaume Adeux, Hana Dvořáková, Anna E-Vojtkó, Ildikó Orbán, Michele Lussu, Javier Puy, et al. 2019. “Colonization Resistance and Establishment Success along Gradients of Functional and Phylogenetic Diversity in Experimental Plant Communities.” Journal of Ecology 107 (5): 2090–2104. 10.1111/1365-2745.13246.

García Criado, Mariana, Isla H. Myers-Smith, Anne D. Bjorkman, Sarah C. Elmendorf, Signe Normand, Peter Aastrup, Rien Aerts, et al. 2025. “Plant Diversity Dynamics over Space and Time in a Warming Arctic.” *Nature*, April, 1–9. 10.1038/s41586-025-08946-8.

Gerhold, Pille, James F. Cahill, Marten Winter, Igor V. Bartish, and Andreas Prinzing. 2015. “Phylogenetic Patterns Are Not Proxies of Community Assembly Mechanisms (they Are Far Better).” Functional Ecology 29 (5): 600–614. 10.1111/1365-2435.12425.

González-Orozco, Carlos E., Laura J. Pollock, Andrew H. Thornhill, Brent D. Mishler, Nunzio Knerr, Shawn W. Laffan, Joseph T. Miller, et al. 2016. “Phylogenetic Approaches Reveal Biodiversity Threats under Climate Change.” Nature Climate Change 6 (12): 1110–14. 10.1038/nclimate3126.

Hähn, Georg J. A., Gabriella Damasceno, Esteban Alvarez-Davila, Isabelle Aubin, Marijn Bauters, Erwin Bergmeier, Idoia Biurrun, et al. 2024. “Global Decoupling of Functional and Phylogenetic Diversity in Plant Communities.” Nature Ecology & Evolution 9 (December):237–48. 10.1038/s41559-024-02589-0.

Harrison, Susan, Ellen I. Damschen, and James B. Grace. 2010. “Ecological Contingency in the Effects of Climatic Warming on Forest Herb Communities.” Proceedings of the National Academy of Sciences of the United States of America 107 (45): 19362–67. 10.1073/pnas.1006823107.

Harte, John, and Rebecca Shaw. 1995. “Shifting Dominance Within a Montane Vegetation Community: Results of a Climate-Warming Experiment.” Science 267 (5199): 876–80. 10.1126/science.267.5199.876.

Huang, Mengjiao, Xiang Liu, Marc W. Cadotte, and Shurong Zhou. 2020. “Functional and Phylogenetic Diversity Explain Different Components of Diversity Effects on Biomass Production.” Oikos 129 (8): 1185–95. 10.1111/oik.07032.

Iler, Amy M., Aldo Compagnoni, David W. Inouye, Jennifer L. Williams, Paul J. CaraDonna, Aaron Anderson, and Tom E. X. Miller. 2019. “Reproductive Losses due to Climate Change-Induced Earlier Flowering Are Not the Primary Threat to Plant Population Viability in a Perennial Herb.” The Journal of Ecology 107 (4): 1931–43. 10.1111/1365-2745.13146.

Inouye, David W. 2008. “Effects of Climate Change on Phenology, Frost Damage, and Floral Abundance of Montane Wildflowers.” Ecology 89 (2): 353–62. 10.1890/06-2128.1.

Kapfer, Jutta, Radim Hédl, Gerald Jurasinski, Martin Kopecký, Fride H. Schei, and John-Arvid Grytnes. 2016. “Resurveying Historical Vegetation Data - Opportunities and Challenges.” Applied Vegetation Science 20 (2): 164–71. 10.1111/avsc.12269.

Kelly, Anne E., and Michael L. Goulden. 2008. “Rapid Shifts in Plant Distribution with Recent Climate Change.” Proceedings of the National Academy of Sciences of the United States of America 105 (33): 11823–26. 10.1073/pnas.0802891105.

Ketola, T., K. Saarinen, and L. Lindström. 2017. “Propagule Pressure Increase and Phylogenetic Diversity Decrease Community’s Susceptibility to Invasion.” BMC Ecology 17 (1): 15. 10.1186/s12898-017-0126-z.

Lambert, Allison M., Abraham J. Miller-Rushing, and David W. Inouye. 2010. “Changes in Snowmelt Date and Summer Precipitation Affect the Flowering Phenology of Erythronium Grandiflorum (glacier Lily; Liliaceae).” American Journal of Botany 97 (9): 1431–37. 10.3732/ajb.1000095.

Langenheim, Jean H. 1962. “Vegetation and Environmental Patterns in the Crested Butte Area, Gunnison County, Colorado.” Ecological Monographs 32 (3): 249–85. 10.2307/1942400.

Langenheim, J. H. 1953. “Plant-Ecological Reconnaissance of the Crested Butte Area.” Minneapolis, MN, USA: University of Minnesota.

Leitão, Rafael P., Jansen Zuanon, Sébastien Villéger, Stephen E. Williams, Christopher Baraloto, Claire Fortunel, Fernando P. Mendonça, and David Mouillot. 2016. “Rare Species Contribute Disproportionately to the Functional Structure of Species Assemblages.” *Proceedings*. Biological Sciences 283 (1828). 10.1098/rspb.2016.0084.

Li, Daijiang, Jesse E. D. Miller, and Susan Harrison. 2019. “Climate Drives Loss of Phylogenetic Diversity in a Grassland Community.” Proceedings of the National Academy of Sciences of the United States of America 116 (40): 19989–94. 10.1073/pnas.1912247116.

López-Rubio, Roberto, David S. Pescador, Adrián Escudero, and Ana M. Sánchez. 2022. “Rainy Years Counteract Negative Effects of Drought on Taxonomic, Functional and Phylogenetic Diversity: Resilience in Annual Plant Communities.” The Journal of Ecology 110 (10): 2308–20. 10.1111/1365-2745.13948.

Maherali, Hafiz, and John N. Klironomos. 2007. “Influence of Phylogeny on Fungal Community Assembly and Ecosystem Functioning.” Science (New York, N.Y.) 316 (5832): 1746–48. 10.1126/science.1143082.

Miller, Jesse E. D., Daijiang Li, Marina LaForgia, and Susan Harrison. 2019. “Functional Diversity Is a Passenger but Not Driver of Drought-related Plant Diversity Losses in Annual Grasslands.” The Journal of Ecology 107 (5): 2033–39. 10.1111/1365-2745.13244.

Miller-Rushing, Abraham J., and David W. Inouye. 2009. “Variation in the Impact of Climate Change on Flowering Phenology and Abundance: An Examination of Two Pairs of Closely Related Wildflower Species.” American Journal of Botany 96 (10): 1821–29. 10.3732/ajb.0800411.

Molina-Venegas, Rafael, Ignacio Ramos-Gutiérrez, and Juan Carlos Moreno-Saiz. 2020. “Phylogenetic Patterns of Extinction Risk in the Endemic Flora of a Mediterranean Hotspot as a Guiding Tool for Preemptive Conservation Actions.” Frontiers in Ecology and Evolution 8 (October):571587. 10.3389/fevo.2020.571587.

Mote, Philip W., Alan F. Hamlet, Martyn P. Clark, and Dennis P. Lettenmaier. 2005. “Declining Mountain Snowpack in Western North America.” Bulletin of the American Meteorological Society 86 (1): 39–50. 10.1175/bams-86-1-39.

Mouillot, David, David R. Bellwood, Christopher Baraloto, Jerome Chave, Rene Galzin, Mireille Harmelin-Vivien, Michel Kulbicki, et al. 2013. “Rare Species Support Vulnerable Functions in High-Diversity Ecosystems.” PLoS Biology 11 (5): e1001569. 10.1371/journal.pbio.1001569.

NOAA. 2014. “Historic Weather Data.” National Centers for Environmental Information. https://www.ncei.noaa.gov/cdo-web/datasets/NORMAL_ANN/locations/ZIP:81224/detail.

Owen, Nisha R., Rikki Gumbs, Claudia L. Gray, and Daniel P. Faith. 2019. “Global Conservation of Phylogenetic Diversity Captures More than Just Functional Diversity.” Nature Communications 10 (1): 859. 10.1038/s41467-019-08600-8.

Padullés Cubino, Josep, Jonathan Lenoir, Daijiang Li, Flavia A. Montaño-Centellas, Javier Retana, Lander Baeten, Markus Bernhardt-Römermann, et al. 2024. “Evaluating Plant Lineage Losses and Gains in Temperate Forest Understories: A Phylogenetic Perspective on Climate Change and Nitrogen Deposition.” New Phytologist 241 (5): 2287–99. 10.1111/nph.19477.

Pagel, Mark. 1999. “Inferring the Historical Patterns of Biological Evolution.” Nature 401 (6756): 877–84. 10.1038/44766.

Palmquist, Kyle A., Daniel R. Schlaepfer, Rachel R. Renne, Stephen C. Torbit, Kevin E. Doherty, Thomas E. Remington, Greg Watson, John B. Bradford, and William K. Lauenroth. 2021. “Divergent Climate Change Effects on Widespread Dryland Plant Communities Driven by Climatic and Ecohydrological Gradients.” Global Change Biology 27 (20): 5169–85. 10.1111/gcb.15776.

Panetta, Anne Marie, Maureen L. Stanton, and John Harte. 2018. “Climate Warming Drives Local Extinction: Evidence from Observation and Experimentation.” Science Advances 4 (2): eaaq1819. 10.1126/sciadv.aaq1819.

Powers, John M., Heather M. Briggs, and Diane R. Campbell. 2025. “Natural Selection on Floral Volatiles and Other Traits Can Change with Snowmelt Timing and Summer Precipitation.” The New Phytologist 245 (1): 332–46. 10.1111/nph.20157.

Powers, John M., Heather M. Briggs, Rachel G. Dickson, Xinyu Li, and Diane R. Campbell. 2022. “Earlier Snow Melt and Reduced Summer Precipitation Alter Floral Traits Important to Pollination.” Global Change Biology 28 (1): 323–39. 10.1111/gcb.15908.

Rudgers, Jennifer A., Stephanie N. Kivlin, Kenneth D. Whitney, Mary V. Price, Nickolas M. Waser, and John Harte. 2014. “Responses of High-Altitude Graminoids and Soil Fungi to 20 Years of Experimental Warming.” Ecology 95 (7): 1918–28. 10.1890/13-1454.1.

Selvi, Federico, Elisa Carrari, and Andrea Coppi. 2016. “Impact of Pine Invasion on the Taxonomic and Phylogenetic Diversity of a Relict Mediterranean Forest Ecosystem.” Forest Ecology and Management 367 (May):1–11. 10.1016/j.foreco.2016.02.013.

Sloat, Lindsey L., Amanda N. Henderson, Christine Lamanna, and Brian J. Enquist. 2015. “The Effect of the Foresummer Drought on Carbon Exchange in Subalpine Meadows.” Ecosystems 18 (3): 533–45. 10.1007/s10021-015-9845-1.

Smith, Stephen A., and Joseph W. Brown. 2018. “Constructing a Broadly Inclusive Seed Plant Phylogeny.” American Journal of Botany 105 (3): 302–14. 10.1002/ajb2.1019.

Stark, Jordan, Rebecca Lehman, Lake Crawford, Brian J. Enquist, and Benjamin Blonder. 2017. “Does Environmental Heterogeneity Drive Functional Trait Variation? A Test in Montane and Alpine Meadows.” *Oikos (Copenhagen*, Denmark) 126 (11): 1650–59. 10.1111/oik.04311.

Thébault, Elisa, Veronika Huber, and Michel Loreau. 2007. “Cascading Extinctions and Ecosystem Functioning: Contrasting Effects of Diversity Depending on Food Web Structure.” Oikos 116 (1): 163–73. 10.1111/j.2006.0030-1299.15007.x.

Veldhuisen, Leah. 2025. “Supplemental Tables for ‘Phylogenetic Patterns over Sixty-Five Years of Vegetation Change across a Montane Elevation Gradient’ by Veldhuisen et Al.” Zenodo. 10.5281/ZENODO.17517272.

Veldhuisen, Leah N., Brian J. Enquist, and Katrina M. Dlugosch. 2023. “Phylogenetic Diversity of Flowering Plants Declines throughout the Growing Season across a Subalpine Elevational Gradient.” bioRxiv. 10.1101/2023.11.06.565878.

Veldhuisen, Leah N., Lorah Seltzer, Jocelyn Navarro, Brian J. Enquist, and Katrina M. Dlugosch. 2025a. “Phylogenetic Diversity and Species Diversity Are Decoupled under Experimental Warming and Cooling in Rocky Mountain Plant Communities.” bioRxivorg. 10.1101/2025.06.11.659139.

Veldhuisen, Leah N., Verónica Zepeda, Brian J. Enquist, and Katrina M. Dlugosch. 2025b. “Rare Species Do Not Disproportionately Contribute to Phylogenetic Diversity in a Subalpine Plant Community.” American Journal of Botany 112 (6): e70061. 10.1002/ajb2.70061.

White, Hannah J., Caroline M. McKeon, Robin J. Pakeman, and Yvonne M. Buckley. 2023. “The Contribution of Geographically Common and Rare Species to the Spatial Distribution of Biodiversity.” Global Ecology and Biogeography: A Journal of Macroecology 32 (10): 1730–47. 10.1111/geb.13734.

Widmer, Stefan, Susanne Riedel, Manuel Babbi, Felix Herzog, Thomas Wohlgemuth, Michael Kessler, and Jürgen Dengler. 2025. “One Century of Change: Stronger Diversity Decline in Lowland than in Mountain Grasslands in Central Europe.” Global Change Biology 31 (10): e70529. 10.1111/gcb.70529.

Willis, Charles G., Brad Ruhfel, Richard B. Primack, Abraham J. Miller-Rushing, and Charles C. Davis. 2008. “Phylogenetic Patterns of Species Loss in Thoreau’s Woods Are Driven by Climate Change.” Proceedings of the National Academy of Sciences 105 (44): 17029–33. 10.1073/pnas.0806446105.

Winter, Marten, Oliver Schweiger, Stefan Klotz, Wolfgang Nentwig, Pavlos Andriopoulos, Margarita Arianoutsou, Corina Basnou, et al. 2009. “Plant Extinctions and Introductions Lead to Phylogenetic and Taxonomic Homogenization of the European Flora.” Proceedings of the National Academy of Sciences of the United States of America 106 (51): 21721–25. 10.1073/pnas.0907088106.

Zorio, Stephanie. 2015. “Analysis of Temporal Change in High Elevation Plant Community Composition, East River Basin, Colorado, USA.” Pocatello, Idaho, USA: Idaho State University.

Zorio, Stephanie Denise, Leah N. Veldhuisen, and Charles F. Williams. 2025. “Langenheim Plant Species Data (1953) and Associated Resurvey Datasets (2014), Gunnison Basin, Colorado, USA.” Environmental Data Initiative. 10.6073/PASTA/9B46B291406E5D44103F78A980BB159F.

Zorio, Stephanie D., Charles F. Williams, and Ken A. Aho. 2016. “Sixty-Five Years of Change in Montane Plant Communities in Western Colorado, U.S.A.” Arctic, Antarctic, and Alpine Research 48 (4): 703–22. 10.1657/AAAR0016-011.

