## Appendix S1 for "Phylogenetic patterns over sixty-five years of vegetation change across a montane elevation gradient"

### Phylogenetic patterns over sixty-five years of vegetation change across a montane elevation gradient

**Authors:** Leah N. Veldhuisen[1,2]* (https://orcid.org/0000-0002-7094-0108), Stephanie D. Zorio[2,3] (https://orcid.org/0000-0001-7847-7667), Charles F. Williams[2] (https://orcid.org/0000-0003-4138-308X), Katrina M. Dlugosch[1] (https://orcid.org/0000-0002-7302-6637)

*Ecosphere*

**SUPPLEMENTAL FIGURES**

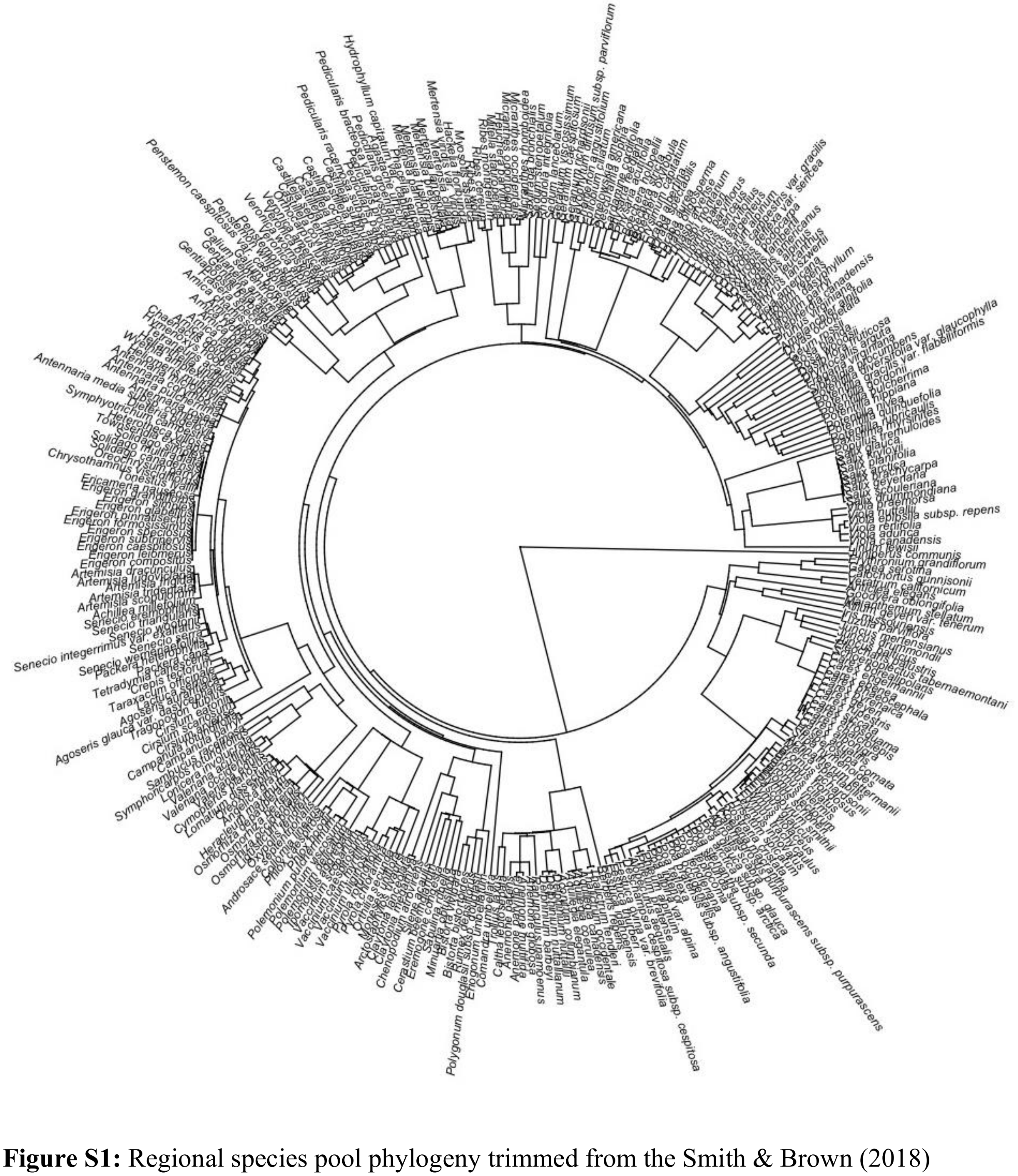


**Figure S1:** Regional species pool phylogeny trimmed from the Smith & Brown (2018) phylogeny.

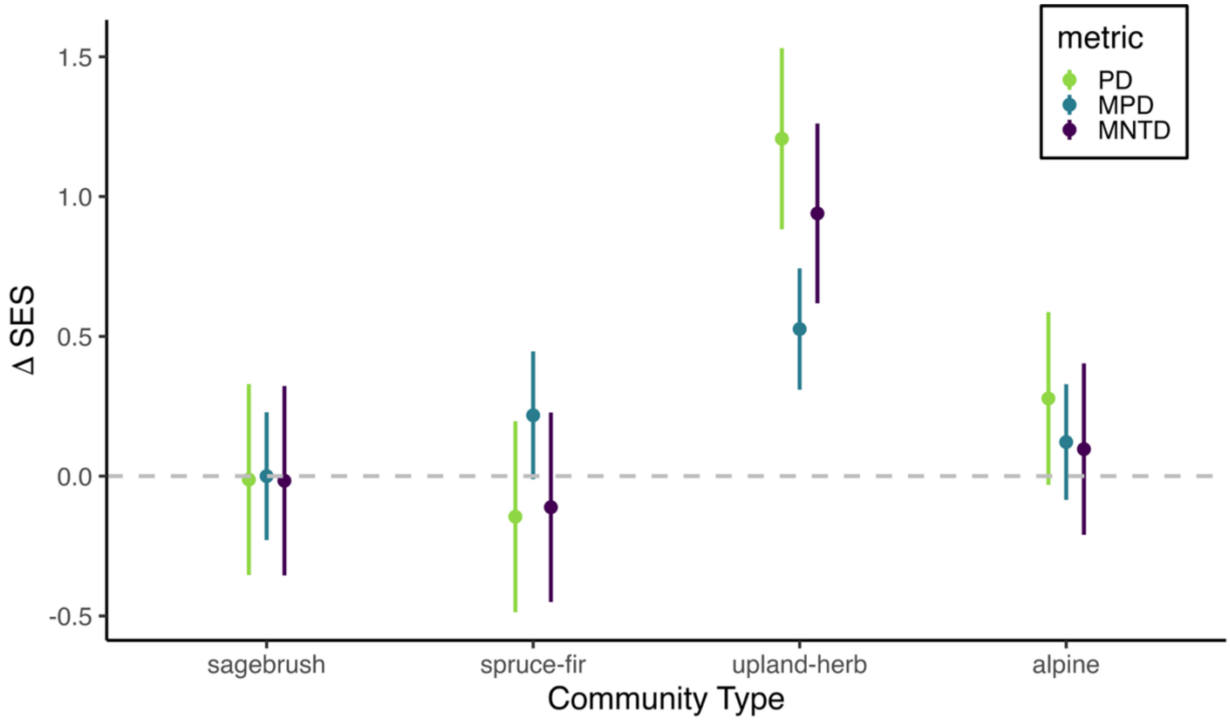


**Figure S2:** Changes in standard effect size for Faith's phylogenetic diversity (green), mean phylogenetic distance (blue) and mean nearest taxon distance (purple) for each community type with the lumped species removed. Points represent means ± SE. Change is statistically significant if the error bars do not cross the grey dashed line, which includes all changes in upland-herb, and none in any other community type.

**References:**

Smith, S. A., and J. W. Brown. 2018. Constructing a broadly inclusive seed plant phylogeny. American journal of botany 105:302–314.
